# STOP1 dominates Arabidopsis tolerance to ammonium over NRT1.1/NPF6.3/CHL1

**DOI:** 10.1101/2024.05.06.592859

**Authors:** Takushi Hachiya, Mako Sakai, Tsuyoshi Nakagawa, Hitoshi Sakakibara

## Abstract

The Arabidopsis nitrate transceptor NITRATE TRANSPORTER 1.1 (NRT1.1/NPF6.3/CHL1) plays significant roles even in the absence of nitrate. The loss-of-function of NRT1.1 alleviates growth suppression and chlorosis under toxic levels of ammonium as the sole nitrogen source. Recently, we reported that acidic stress-inducible genes are downregulated by *NRT1*.*1* deficiency under ammonium toxicity conditions, implying that NRT1.1 may exacerbate ammonium-dependent acidic stress. The transcription factor SENSITIVE TO PROTON RHIZOTOXICITY 1 (STOP1) enhances Arabidopsis tolerance to ammonium and acidic stresses. Furthermore, STOP1 directly activates *NRT1*.*1* transcription. These previous findings prompted us to analyze the association between NRT1.1 and STOP1 on ammonium tolerance. Here, we show that ammonium-inducible STOP1 plays a dominant role over NRT1.1 in ammonium tolerance and expression of direct target genes of STOP1. We present a novel scheme in which NRT1.1 and STOP1 jointly regulate common target(s) associated with ammonium tolerance.

## 1. Introduction

Nitrate serves not only as a substrate for nitrogen (N) assimilation but also as a signaling molecule. The Arabidopsis NITRATE TRANSPORTER 1.1 (NRT1.1/NPF6.3/CHL1) acts as a nitrate sensor and orchestrates various plant responses to the nitrate signal, including gene expression and root system development (Bouguyon et al. 2015; Ho et al. 2009). Apart from its nitrate-dependent functions, NRT1.1 exhibits significant roles in the absence of nitrate (Hachiya et al. 2024). Previous studies have shown that the loss-of-function of NRT1.1 alleviates growth suppression and chlorosis under toxic levels of ammonium as the sole N source, i.e., ammonium toxicity (Hachiya et al. 2011; Jian et al. 2018; Liu et al. 2020). Recently, we reported that in early seedlings grown under 10 mM ammonium, the genes induced by acidic stress are downregulated in *NRT1*.*1*-deficient mutants relative to the wild-type, whereas those repressed by acidic stress are upregulated in the mutants (Hachiya et al. 2024). Moreover, an increase in medium pH promotes shoot growth of the wild-type more intensely than that of the *NRT1*.*1*-deficient mutant under 10 mM ammonium. Given that acidic stress is one of the primary causes of ammonium toxicity (Hachiya et al. 2021), NRT1.1 may exacerbate the ammonium-dependent acidic stress.

In *Arabidopsis*, the zinc finger transcription factor SENSITIVE TO PROTON RHIZOTOXICITY 1 (STOP1) positively regulates genes involved in the tolerance to ammonium and acidic stresses, as well as abiotic stresses, including aluminum, drought, salt, low phosphate, and low oxygen (Balzergue et al. 2017; Enomoto et al. 2019; Hachiya et al. 2021; Sadhukhan et al. 2019; Sawaki et al. 2009; Tian et al. 2021). Furthermore, STOP1 directly activates *NRT1*.*1* expression under an acidic rhizosphere containing nitrate as the primary N source (Ye et al. 2021). These findings prompted us to analyze the association between NRT1.1 and STOP1 under toxic levels of ammonium as the sole N source. Here, we show that STOP1 dominates ammonium tolerance and expression of direct target genes of STOP1 over NRT1.1.

## 2. Materials and Methods

### 2.1. Plant materials and growth conditions

*Arabidopsis thaliana* accession Columbia (Col-0) was used as a control. T-DNA insertion mutants *nrt1*.*1* (SALK_097431, Hachiya et al. 2011), *stop1*^*KO*^ (SALK_114108, Sawaki et al. 2007), and gamma ray-mutagenized mutant c*hl1-5* (Tsay et al. 1993) were obtained from the European Arabidopsis Stock Center. *stop1*^*KO*^ was backcrossed twice with Col-0 before use. The line harboring *pSTOP1:GFP-STOP1* in the *stop1*^*KO*^ background (Balzergue et al. 2017) was provided by Dr. Thierry Desnos (Aix-Marseille University, France).

The seeds were surface sterilized, placed in plastic Petri dishes (diameter: 90 mm; depth: 20 mm; Iwaki, Tokyo, Japan), and grown on a solid medium containing 30 mL of Murashige and Skoog (MS) salts without N sources supplemented with 4.7 mM MES-KOH (pH 5.7), 2% (w/v) sucrose, and 0.25% (w/v) gellan gum (Wako, Osaka, Japan) with or without 100 nM indole-3-acetic acid (IAA). Two different N and K sources were used: 10 mM KNO3 (10 mM NO3− condition) or 5 mM (NH4)2SO_4_ with 10 mM KCl (10 mM NH_4+_ condition). The plants were kept in the dark at 4°C for 3 days, and then grown horizontally under a photosynthetic photon flux density of 100–130 μmol m^−2^ s^−1^ (16-h light/8-h dark cycle) at 23°C.

### 2.2. Extraction of RNA

Frozen samples were ground using TissueLyser II (Qiagen, Tokyo, Japan) using zirconia beads. Total RNA was extracted using an RNeasy Plant Mini Kit (Qiagen), following the manufacturer’s instructions.

### 2.3. Quantitative reverse transcription polymerase chain reaction

Reverse transcription (RT) was conducted using ReverTra Ace qPCR RT Master Mix with gDNA Remover (Toyobo, Tokyo, Japan), following the manufacturer’s instructions. The obtained cDNA was diluted 10-fold with distilled water for quantitative PCR (qPCR). Transcript levels were evaluated using StepOnePlus Real-Time PCR System (Thermo Fisher Scientific, Waltham, MA, USA). The diluted cDNA (2 µL) was amplified in the presence of 10 µL of KAPA SYBR FAST qPCR Kit (Nippon Genetics, Tokyo, Japan), 0.4 µL of specific primers (0.2 µM final concentration), and 7.2 µL of sterile water. Relative transcript levels were calculated using the comparative cycle threshold method with *ACTIN3* as an internal standard. Primer sequences are shown in Supplementary Table 1.

### 2.4. Observation of fluorescent signal

Confocal imaging was performed using a Leica SP5 Confocal Microscope (Leica Microsystems, Wetzlar, Germany). GFP-STOP1 was excited with a 488-nm Ar laser, and emission was detected from 500–530 nm using a Leica 25× objective (HCX IRAPO L 25×/0.95 WATER).

## 3. Results and Discussion

### 3.1. STOP1 dominates ammonium tolerance over NRT1.1

To analyze the genetic relationship between *NRT1*.*1* and *STOP1, chl1-5* and *nrt1*.*1* were crossed with *stop1* ^*KO*^, and their double homozygous mutants were obtained (Figure 1a). The growth of Col-0, *chl1-5, nrt1*.*1, stop1*^*KO*^, *chl1-5 stop1*^*KO*^, and *nrt1*.*1 stop1*^*KO*^ under 10 mM ammonium supplemented with 100 nM IAA was compared. It should be noted that appropriate IAA concentrations stabilize the ammonium tolerance of *NRT1*.*1*-deficient mutants (Hachiya and Noguchi 2011). The shoots of *chl1-5* and *nrt1*.*1* were larger and greener than those of Col-0 (Figure 1b). The shoot growth enhancement in *NRT1*.*1*-deficient mutants was almost completely suppressed by *STOP1* deficiency. Shoot fresh weights of *chl1-5* and *nrt1*.*1* were much larger than those of Col-0, whereas those of *stop1*^*KO*^ were slightly smaller (Figure 1c). Fresh weights of the shoots of *chl1-5 stop1*^*KO*^ and *nrt1*.*1 stop1*^*KO*^ were comparable to those of *stop1*^*KO*^. Also, under 10 mM ammonium without IAA, the shoot fresh weights were increased by *NRT1*.*1* deficiency, but they were reduced by *STOP1* deficiency to a level comparable to those of *stop1*^*KO*^ (Figure 1d). Under 10 mM nitrate without IAA, the shoot fresh weights of *NRT1*.*1*-deficient mutants were significantly smaller than those of other lines with or without *STOP1*. Under both N conditions, the fresh weights of the root showed a similar tendency to those of the shoot. Thus, our genetic analysis indicates that STOP1 is epistatic to NRT1.1 regarding ammonium tolerance.

**Figure 1.**
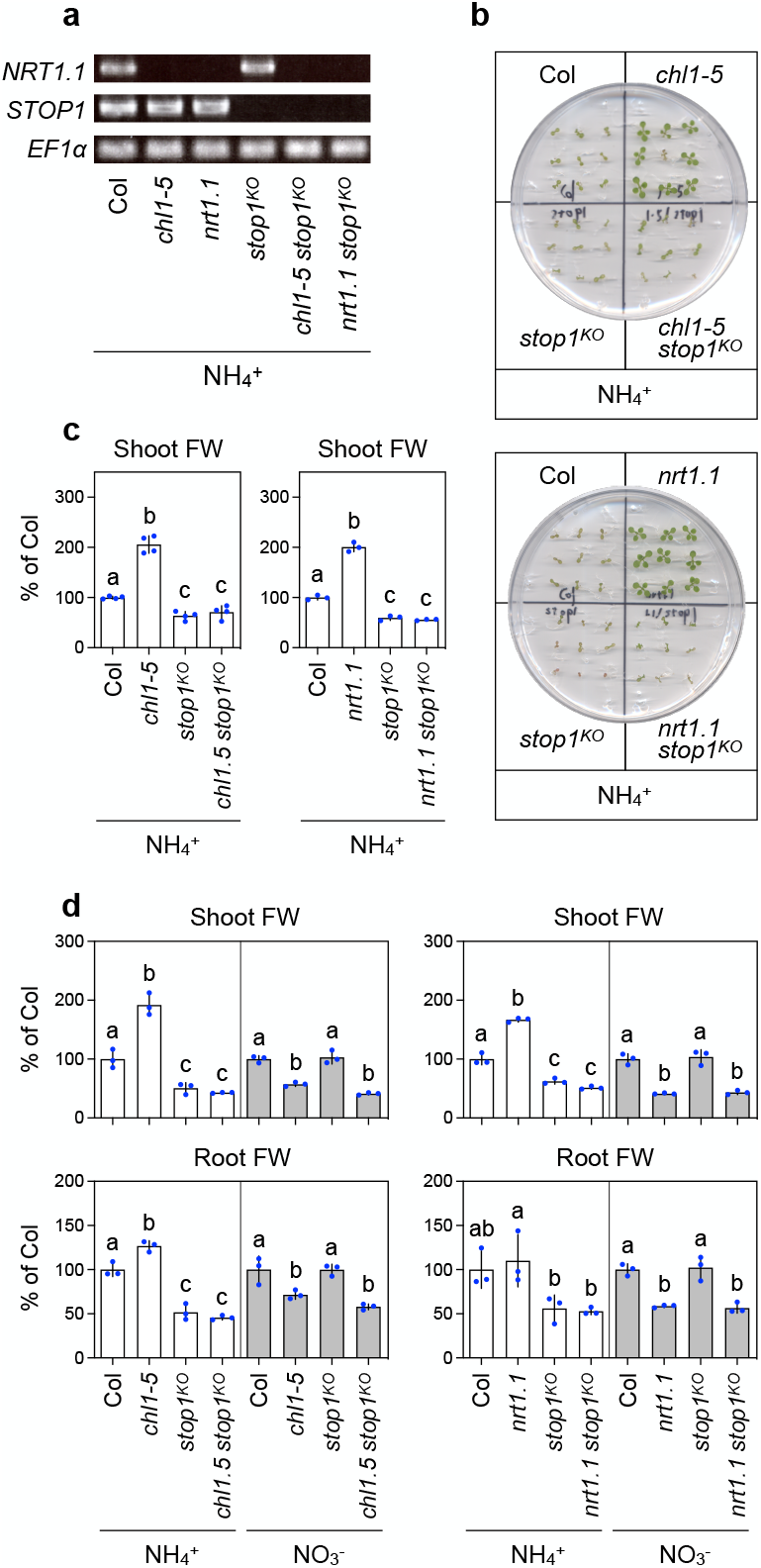
*STOP1* deficiency suppresses enhanced ammonium tolerance in *NRT1*.*1*-deficient mutants. (a) RT-PCR and agarose gel electrophoresis for *NRT1*.*1, STOP1*, and *EF1α* using five-day-old seedlings of Col-0, *chl1-5, nrt1*.*1, stop1*^*KO*^, *chl1-5 stop1*^*KO*^, and *nrt1*.*1 stop1*^*KO*^ grown under 10 mM ammonium. (b,c) Representative photographs (b) and fresh weights (c) of shoots from 11-day-old Col-0, *chl1-5, nrt1*.*1, stop1*^*KO*^, *chl1-5 stop1*^*KO*^, and *nrt1*.*1 stop1*^*KO*^ seedlings grown under 10 mM ammonium with 100 nM IAA (using 100 µM IAA (in dimethyl sulfoxide) stock solution). (d) Fresh weights of shoots and roots from 11-day-old Col-0, *chl1-5, nrt1*.*1, stop1*^*KO*^, *chl1-5 stop1*^*KO*^, and *nrt1*.*1 stop1*^*KO*^ seedlings grown under 10 mM ammonium without IAA. (b-d) In one dish, nine seedlings of each line were regarded as a single biological replicate (Mean ± SD; n = 3–4). Different lowercase letters indicate significant differences evaluated by the Tukey–Kramer multiple comparison test conducted at a significance level of *P* <0.05.

### 3.2. NRT1.1 does not downregulate STOP1/STOP1 and STOP1 dominates expression of direct target genes of STOP1 over NRT1.1

The above growth analysis suggests that NRT1.1 could act upstream of STOP1 and downregulate *STOP1*/STOP1 activities under ammonium. To verify this, we checked the transcript levels of known direct target genes of STOP1, i.e., *ALUMINUM-ACTIVATED MALATE TRANSPORTER 1* (*ALMT1*), *CALCINEURIN B-LIKE INTERACTING PROTEIN KINASE 23* (*CIPK23*), *GLUTAMATE DEHYDROGENASE 2* (*GDH2*), and *SENSITIVE TO PROTON RHIZOTOXICITY 2* (*STOP2*) (Balzergue et al. 2017; Enomoto et al. 2019; Sadhukhan et al. 2019, Tokizawa et al. 2021) in the absence of IAA. In five-day-old Col-0 shoots and roots, the expression of *ALMT1, CIPK23, GDH2*, and *STOP2* was significantly higher under 10 mM ammonium than under 10 mM nitrate (Figure 2a). However, this ammonium-dependent transcriptional induction was consistently decreased by *NRT1*.*1* deficiency (Figure 2a). Also, *STOP1* expression in shoots and roots was not significantly increased by *NRT1*.*1* deficiency under ammonium (Figure 2a). Moreover, our microarray analysis using three- and five-day-old seedlings under 10 mM ammonium without IAA (Hachiya et al. 2024) revealed that *NRT1*.*1* deficiency did not generally induce genes whose expression is upregulated via STOP1 under acidic pH (Sawaki et al. 2009) (Figure 2b and Supplementary Table S2). It is widely accepted that the F-box protein REGULATION OF ATALMT1 EXPRESSION 1 (RAE1) interacts with STOP1 and promotes its degradation via the ubiquitin-26S proteasome pathway (Zhang et al. 2019), i.e., the post-translational regulation of STOP1 activity. Recent studies have shown that aluminum stress induces phosphorylation of STOP1 by MAP KINASE 4 and oxidative modification of RAE1 by reactive oxygen species (Zhou et al. 2023; Ding et al. 2024). These modifications inhibit RAE1-dependent proteolysis of STOP1, thereby enhancing aluminum tolerance. Thus, to analyze the effect of NRT1.1 on STOP1 expression, *chl1-5* was crossed with *pSTOP1:GFP-STOP1* line in the *stop1*^*KO*^ background (Balzergue et al. 2017) and produced the *pSTOP1:GFP-STOP1* line in the *chl1-5 stop1*^*KO*^ background. The GFP-STOP1-derived fluorescence signal was promoted in the roots under 10 mM ammonium compared to 10 mM nitrate (Figure 2c), similar to a previous result (Tian et al. 2021). However, the GFP signal was little changed by *NRT1*.*1* deficiency. Meanwhile, in five-day-old seedlings grown under 10 mM ammonium, the transcript levels of *ALMT1, CIPK23*, and *GDH2* were similar among *stop1*^*KO*^, *chl1-5 stop1*^*KO*^, and *nrt1*.*1 stop1*^*KO*^, all of which were lower than those observed in *chl1-5* and *nrt1*.*1* (Figure 2d). These observations suggest that NRT1.1 does not downregulate *STOP1*/STOP1 activities and that STOP1 dominates expression of direct target genes of STOP1 over NRT1.1.

**Figure 2.**
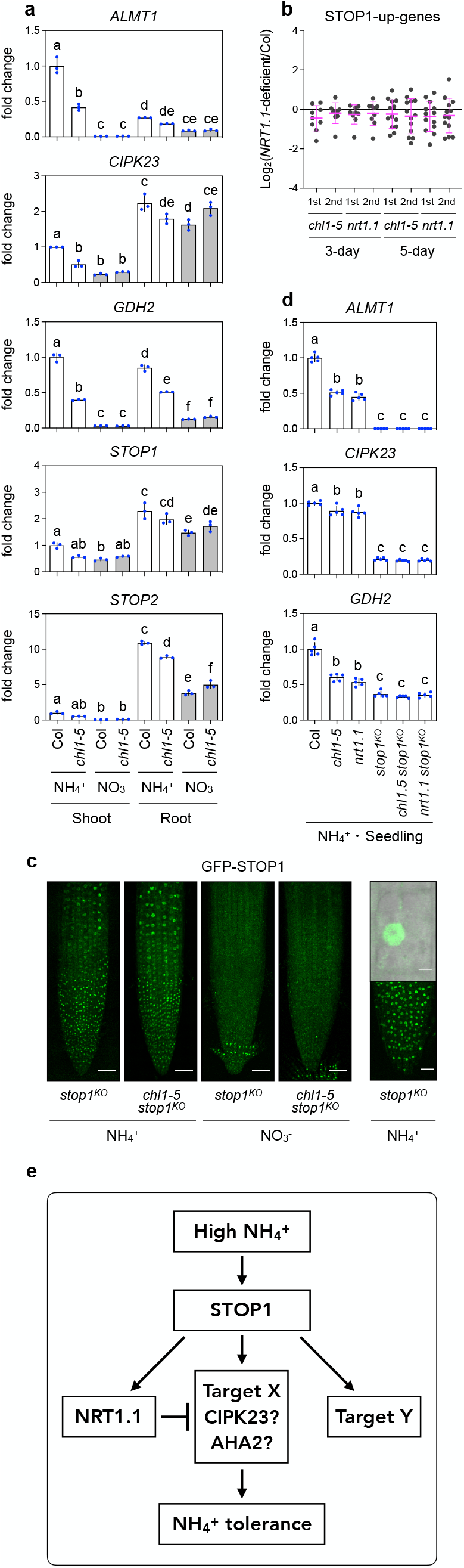
NRT1.1 does not downregulate *STOP1*/STOP1 and STOP1 dominates expression of direct target genes of STOP1 over NRT1.1. (a) Relative transcript levels of *ALMT1, CIPK23, GDH2, STOP1*, and *STOP2* in shoots and roots of five-day-old seedlings from Col-0 and *chl1-5* grown under 10 mM ammonium or 10 mM nitrate conditions without IAA. (b) Comparisons of the expression of STOP1-upregulated genes under acidic pH (Sawaki et al. 2009) between Col-0 and *NRT1*.*1*-deficient mutants (*chl1-5, nrt1*.*1*) grown under 10 mM ammonium without IAA based on the microarray results (Hachiya et al. 2024). Changes in gene expression levels were shown as logarithms to base 2 of fold change (Mean ± SD). (c) GFP-STOP1 signals from the primary root tips of five-day-old seedlings grown under 10 mM ammonium or 10 mM nitrate without IAA. The left four images and lower right image were generated by the maximum intensity projection of multiple slices of the range where fluorescence is clearly observed. The single cell image in the upper right is a composite of bright and fluorescent images. Scale bars: 50 µm (left), 5 µm (upper right), and 20 µm (lower right). (d) Relative transcript levels of *ALMT1, CIPK23*, and *GDH2* in five-day-old seedlings from Col-0, *chl1-5, nrt1*.*1, stop1*^*KO*^, *chl1-5 stop1*^*KO*^, and *nrt1*.*1 stop1*^*KO*^ grown under 10 mM ammonium without IAA. (a, d) Thirty-seven seedlings from one plate were regarded as a single biological replicate (Mean ± SD; n = 3–5). (e) A possible scheme for ammonium tolerance regulation by NRT1.1 and STOP1.

### 3.3 Possible scheme for regulation of ammonium tolerance by NRT1.1 and STOP1

Ye et al. (2021) demonstrated that, under acidic conditions with nitrate as the primary N source, STOP1 directly upregulates *NRT1*.*1* expression in the root, which facilitates co-uptake of proton and nitrate, elevating rhizosphere pH to a favorable level for *Arabidopsis*: therefore, NRT1.1 was epistatic to STOP1 on root nitrate uptake and root elongation. Meanwhile, the present study showed that, under toxic levels of ammonium without nitrate, ammonium-inducible STOP1 dominates ammonium tolerance and target gene expression of STOP1 over NRT1.1 (Figure 1 and 2). Moreover, we found that ammonium-dependent transcriptional induction of shoot *NRT1*.*1* was completely suppressed by *STOP1* deficiency (Supplementary Figure 1), supporting that *NRT1*.*1* is under the control of STOP1. To reconcile these observations, a possible scheme for regulation of ammonium tolerance by NRT1.1 and STOP1 is presented (Figure 2e). The STOP1 upregulated by ammonium induces the expression of *NRT1*.*1* and other target gene(s). The function(s) of some target gene product(s) (i.e., Target X in Figure 2e) could enhance ammonium tolerance but be suppressed by NRT1.1, possibly via protein–protein interaction(s).

In *A. thaliana*, STOP1 directly induces *CIPK23* expression under toxic levels of ammonium as the sole N source (Tian et al. 2021). CIPK23 inhibits ammonium transport via phosphorylation of high-affinity AMMONIUM TRANSPORTER 1;1 (AMT1;1) and 1;2 (AMT1;2) (Straub et al. 2017). Since the loss-of-function of *CIPK23* causes excessive ammonium uptake and thus ammonium hypersensitivity (Hachiya et al. 2024; Straub et al. 2017), STOP1 improves ammonium tolerance by upregulating *CIPK23* (Tian et al. 2021). Notably, under low nitrate conditions, CIPK23 physically interacts with NRT1.1 and phosphorylates it, increasing the affinity of NRT1.1 for nitrate (Ho et al. 2009). It remains unclear whether NRT1.1 reduces ammonium tolerance through its interaction with CIPK23 as a candidate of Target X (Figure 2e). Meanwhile, a recent study reported that STOP1 indirectly activates the ARABIDOPSIS PLASMA-MEMBRANE H^+^-ATPase 2 (AHA2) by upregulating *SMALL AUXIN-UP RNA 55* (*SAUR55*) whose translational product prevents PP2C.D2/5, negative regulators of AHA2 (Agrahari et al. 2023). The plant Cistrome database implies that STOP1 could directly bind to the promoter of *AHA2*. By contrast, NRT1.1 and receptor-kinase QIAN SHOU KINASE 1 (QSK1) inactivate the AHA2 via phosphorylation on S899 under low nitrate (Zhu et al. 2024). Hence, STOP1 and NRT1.1 may affect ammonium-derived acidic stress via AHA2 in opposite directions and therefore, AHA2 could also be a possible candidate for Target X in Figure 2e. It would be valuable to explore the common target(s) of NRT1.1 and STOP1 for better understanding ammonium toxicity.

## Acknowledgments

We would like to thank Dr. Thierry Desnos (Aix-Marseille University) for providing the *Arabidopsis* seeds, Dr. Alain Gojon for valuable discussion, and Mr. Shun Ohno for supporting experiments. The authors would like to thank Enago (www.enago.jp) for the English language review.

## Disclosure statement

The authors report there are no competing interests to declare.

## Funding

This study was supported by the Grants-in-Aid from the Ministry of Education, Culture, Sports, Science and Technology, Japan under Grant [17K15237/20K05771/23K04978], the Inamori Foundation, the Agropolis Foundation Grant [1502-405], and the Nagase Science and Technology Foundation, and the Yanmar Joint Research to TH.

## Notes

### Competing Interest Statement

The authors have declared no competing interest.

### Summary of Updates

(1) Addition of Supplementary Figure 1, citations, and the related descriptions. (2) Replacing the order of the figures in Figure 2. (3) Correction of mistakes.

