## Supplementary Figure 1 for "STOP1 dominates Arabidopsis tolerance to ammonium over NRT1.1/NPF6.3/CHL1"

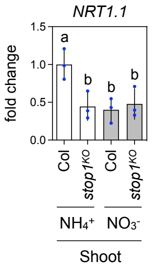


**Figure S1.** STOP1 induces shoot expression of *NRT1.1* under ammonium.

Relative transcript levels of *NRT1.1* in shoots of five-day-old seedlings from Col-0 and *stop1*$^{KO}$ grown under 10 mM ammonium or 10 mM nitrate conditions without IAA. Thirty-seven seedlings from one plate were regarded as a single biological replicate (Mean ± SD; n = 3).
